# Selective Affimers Recognize BCL-2 Family Proteins Through Non-Canonical Structural Motifs

**DOI:** 10.1101/651364

**Authors:** Jennifer A. Miles, Fruzsina Hobor, James Taylor, Christian Tiede, Philip R. Rowell, Chi H. Trinh, Brian Jackson, Fatima Nadat, Hannah F. Kyle, Basile I. M. Wicky, Jane Clarke, Darren C. Tomlinson, Andrew J. Wilson, Thomas A. Edwards

**Affiliations:** School of Molecular and Cellular Biology, Faculty of Biological Sciences, University of Leeds, Woodhouse Lane, Leeds LS2 9JT, UK; Astbury Centre For Structural Molecular Biology, University of Leeds, Woodhouse Lane, Leeds LS2 9JT, UK; School of Chemistry, University of Leeds, Woodhouse Lane, Leeds LS2 9JT, UK; Protein Production Facility, Faculty of Biological Sciences, University of Leeds, Woodhouse Lane, Leeds LS2 9JT, UK; Department of Chemistry, University of Cambridge, Lensfield Road, Cambridge CB2 1EW

## Abstract

The BCL-2 family is a challenging set of proteins to target selectively due to sequence and structural homologies across the family. Selective ligands for the BCL-2 family regulators of apoptosis are desirable as probes to understand cell biology and apoptotic signalling pathways, and as starting points for inhibitor design. We have used phage display to isolate Affimer reagents (non-antibody binding proteins based on a conserved scaffold) to identify ligands for MCL-1, BCL-x_L_, BCL-2, BAK and BAX, then used multiple biophysical characterisation methods to probe the interactions. We established that purified Affimers elicit selective and potent recognition of their target BCL-2 protein. For anti-apoptotic targets, competitive inhibition of their canonical protein-protein interactions is demonstrated. Co-crystal structures reveal an unprecedented mode of molecular recognition; where a BH3 helix is normally bound, flexible loops from the Affimer dock into the BH3 binding cleft. Moreover, the Affimers induce a change in the target proteins towards a desirable drug bound like conformation. These results indicate Affimers can be used as alternative templates to inspire design of selective BCL-2 family modulators, and provide proof-of-concept for the elaboration of selective non-antibody binding reagents for use in cell-biology applications.

## Introduction

A central challenge in life sciences research is to identify modulators of protein-protein interactions (PPIs).^1,2^ Such modulators represent probes with which to uncover new understanding of structural and cellular biology, as well as starting points for drug discovery. The BCL-2 family of PPIs are an important class of α-helix mediated interaction that control the intrinsic apoptosis pathway.^3^ Their critical role in apoptosis has prompted efforts to identify modulators so as to facilitate greater understanding of both BCL-2 family signaling and drug discovery.^4–8^ Moreover differing selectivities and specificities amongst BCL-2 family member interactions^9,10^ render the family an outstanding model system to elaborate novel generic chemical and biological approaches for protein-protein interaction modulation.^11,12^ BCL-2 family proteins can be identified through their BH (BCL-Homology) domains and may be categorized within 3 specific sub-groups (Fig. 1a). Pro-apoptotic (or executioner) proteins such as BAK and BAX activate apoptosis through pore formation in the mitochondrial membrane; anti-apoptotic proteins including BCL-2, MCL-1 and BCL-x_L_, sequester pro-apoptotic members to prevent cell death; and a group of regulatory proteins which bind to other BCL-2 members (including BIM, BID, BAD, NOXA and PUMA), mediate initiation of apoptosis. In all cases binding between BCL-2 family members occurs through the BH3 homology domain of one protein, which forms an α-helix upon binding and docks into a complementary cleft on its partner (Fig. 1b). The BH3 ligand exploits conserved hydrophobic residues in positions *i*, *i* + 4, *i* +7 and *i* +11 together with a conserved aspartic acid (at *i* + 9) to achieve high affinity interaction with the BH3 cleft (Fig. 1c). *In silico* and experimental approaches have been used to identify selective sequences for individual BCL-2 family members.^13–16^ Multiple studies have endeavoured to identify chemotypes which mimic the BH3 domains so as to orthosterically inhibit BCL-2/BH3 PPIs including: constrained peptides,^17–23^ peptidomimetics,^24–27^ small molecules^28–31^ and miniature proteins (identified with assistance from biological selection).^32,33^ We have used a previously described *Affimer* library^34–39^ to identify potent ligands for MCL-1, BCL-x_L_, BCL-2, BAK and BAX and selective inhibitors of MCL-1 and BCL-x_L_ interactions with cognate BH3 partners. Our aim was not only to identify high affinity binders, but also to then screen for subsets that would inhibit PPIs, provide multiple sequences for motif identification, and to use those amenable to structural studies to understand the mode of binding. Affimer reagents belong to an emerging class of non-antibody based protein scaffolds which include, Monobodies, Darpins, Affibodies and others,^40–44^ which may offer advantages in therapeutic and diagnostic settings associated with improved solubility, purification, expression and stability. Affimer reagents are based on either a human scaffold,^45^ or a phytocystatin scaffold (Fig. 1d) which has been optimized by homology.^39^ Both show high thermal stability and achieve molecular recognition through one or two variable regions (VRs) of between six and twelve amino acid residues. Multiple large libraries of Affimers have been established permitting biological selection of optimized binding reagents through randomization within each of the VRs.^39,46^ These reagents provide access to distinct compositional and conformational peptide diversity compared to natural biological peptides, and can be identified *via* the power of genetics rather than synthetic chemistry.

**Figure 1.**
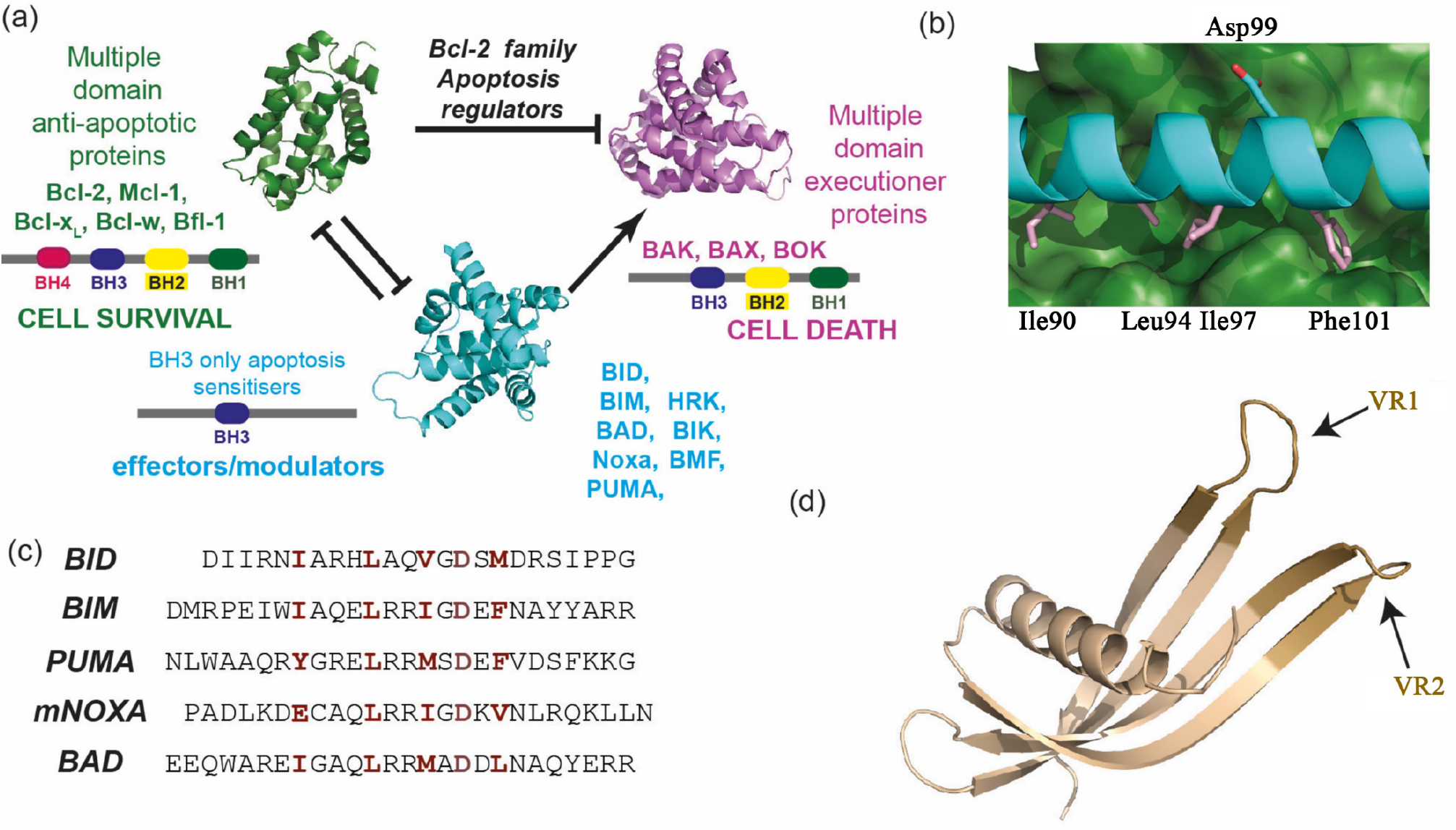
BCL-2 family structure and function (a) schematic annotating BCL-2 family member role in apoptosis (b) BCL-x_L_/BIM co-crystal structure (PDB ID: 1PQ1 highlighting key residues for binding (labelled above and below) (c) sequences of BH3 modulators (annotating key residues required for BH3 cleft affinity in dark red) (d) crystal structure of an Affimer highlighting VRs (dark brown) where amino acid variation is possible (PDB ID: 5A0O) VR = variable region.^38^

## Results

### Isolating Affimers

Following expression using established methods (see ESI), MCL-1, BCL-x_L_, BCL-2, BAK and BAX were biotinylated and immobilised on plates, over which the library of Affimers was panned in order to isolate high-affinity binders (see methods). Phage ELISA was then used to identify clones that bind selectively for further analysis. Following this screening, candidate Affimer reagents were sequenced resulting in twelve unique sequences with affinity for MCL-1 (from 24 clones), eleven for BCL-x_L_ (21 clones), four for BCL-2 (20 clones), five for BAX (31 clones) and four for BAK (24 clones). Tables S1 and S2 (see ESI) indicates the identified sequences and frequency.

### Binding Analysis of Affimers

Small scale expression of the Affimers then allowed preliminary biophysical/biochemical analyses. For MCL-1 and BCL-x_L_, single concentration fluorescence anisotropy (FA) competition assays (Fig. 2 a,b) against BCL-x_L_ /BAK or MCL-1/NOXA-B (using competitor Affimer at 1 µM) were used to identify Affimers that inhibit cognate BH3 binding. Inhibition was compared to positive controls BAK and ABT-737 for BCL-x_L_ and NOXA-B for MCL-1 with the peptide activity defined as 100% inhibition (note that ABT-737 is more active than BAK therefore achieves 150% inhibition). From these assays, three BCL-x_L_ Affimers were identified with significant inhibitory potency, (**BCL-x_L_-AF6**, **BCL-x_L_-AF7** and **BCL-x_L_-AF10**) and two MCL-1 Affimers (**MCL-1-AF1** and **MCL-1-AF11**). We did not have an established competition assay for BAK and BAX. BAK, BAX and BCL-2 Affimers were purified by size exclusion chromatography, then confirmation of correct Affimer folding was obtained through circular dichroism (CD, see ESI, Fig. S1). Binding ELISA using a primary antibody for the His tag on the Affimer and secondary HRP antibody established potent and selective interaction between the selected Affimer and BCL-2 targets (Fig. 2 c,d and Fig. S2). **BCL-2-AF1** to **BCL-2-AF3** and **BAK-AF1** to **BAK-AF4** were confirmed as genuine binders, selective for their targets, but no BAX Affimers were successfully confirmed from the ELISA analyses.

**Figure 2.**
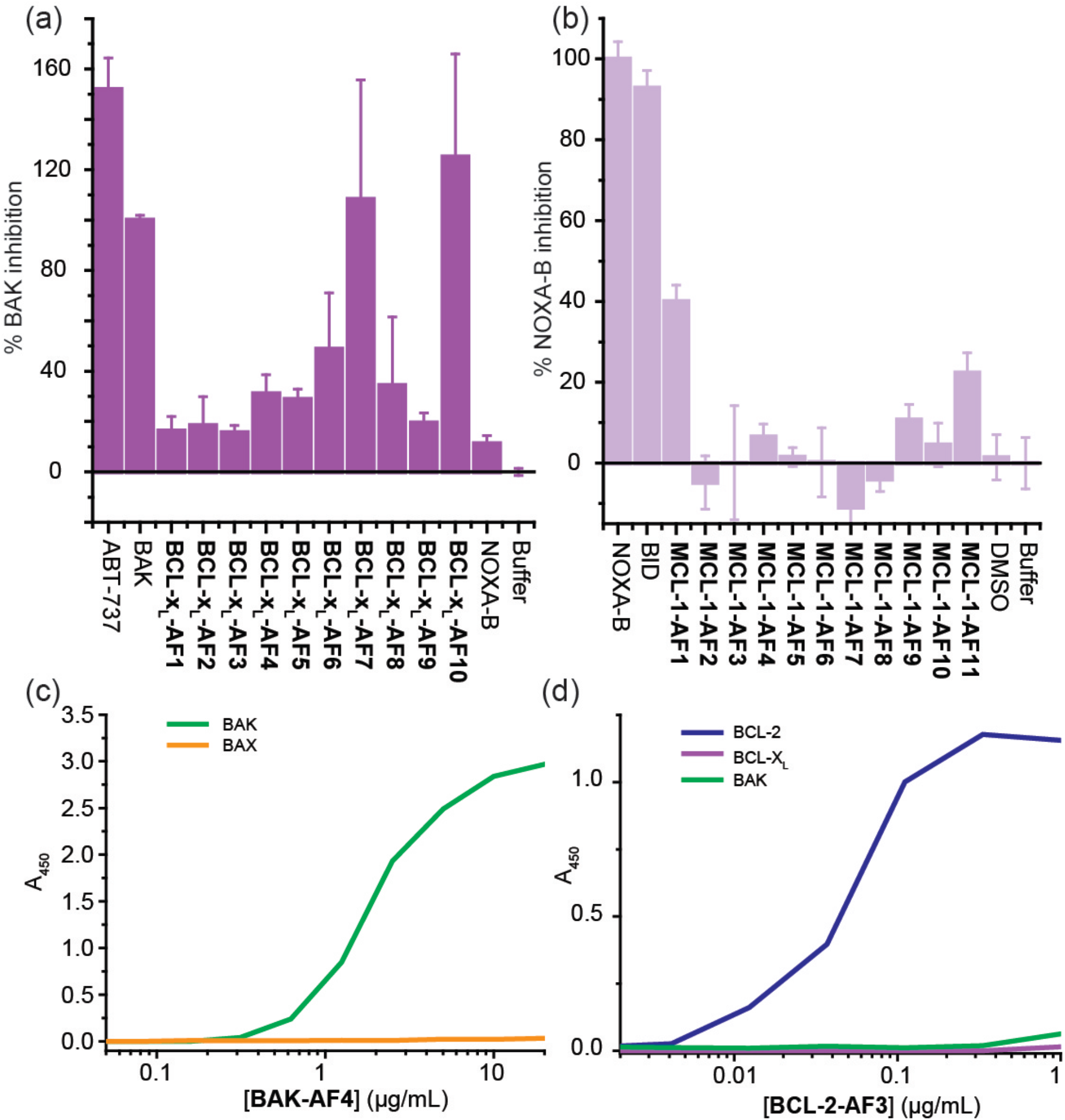
Binding analysis and selectivity of BCL-2 family binding Affimers. Single point screening of (a) BCL-x_L_ binding Affimers (1µM) for competitive inhibition of BH3 binding and (b) MCL-1 binding Affimers (1 µM); binding ELISA for (c) **BAK-AF4** (d) **BCL-2-AF3,**

### Biophysical Analysis of Affimers

Larger scale expression in *E. coli* of the five Affimers identified from single point FA competition allowed the purified proteins to be tested in full dose response fluorescence anisotropy competition assays against their target (Fig. 3 a,b). **BCL-x_L_-AF10** showed problems during purification so was not further characterized. Both **BCL-x_L_-AF6** (IC_50_ = 448 ± 53 nM) and **BCL-x_L_-AF7** (IC_50_ = 393 ±54 nM) were shown to act as sub µM inhibitors of the BCL-x_L_/BAK interaction (Fig. 3a) but were ineffective in inhibiting the MCL-1/NOXA-B interaction. Similarly, the Affimers selected for MCL-1 binding were shown to act as low µM inhibitors of their target interaction (**MCL-1-AF1** IC_50_ = 2.1 ± 0.2 μM; **MCL-1-AF11** IC_50_ = 3.2 ± 0.4 μM) but did not inhibit BCL-x_L_ /BAK, (Fig. 3b), demonstrating selectivity.

**Figure 3.**
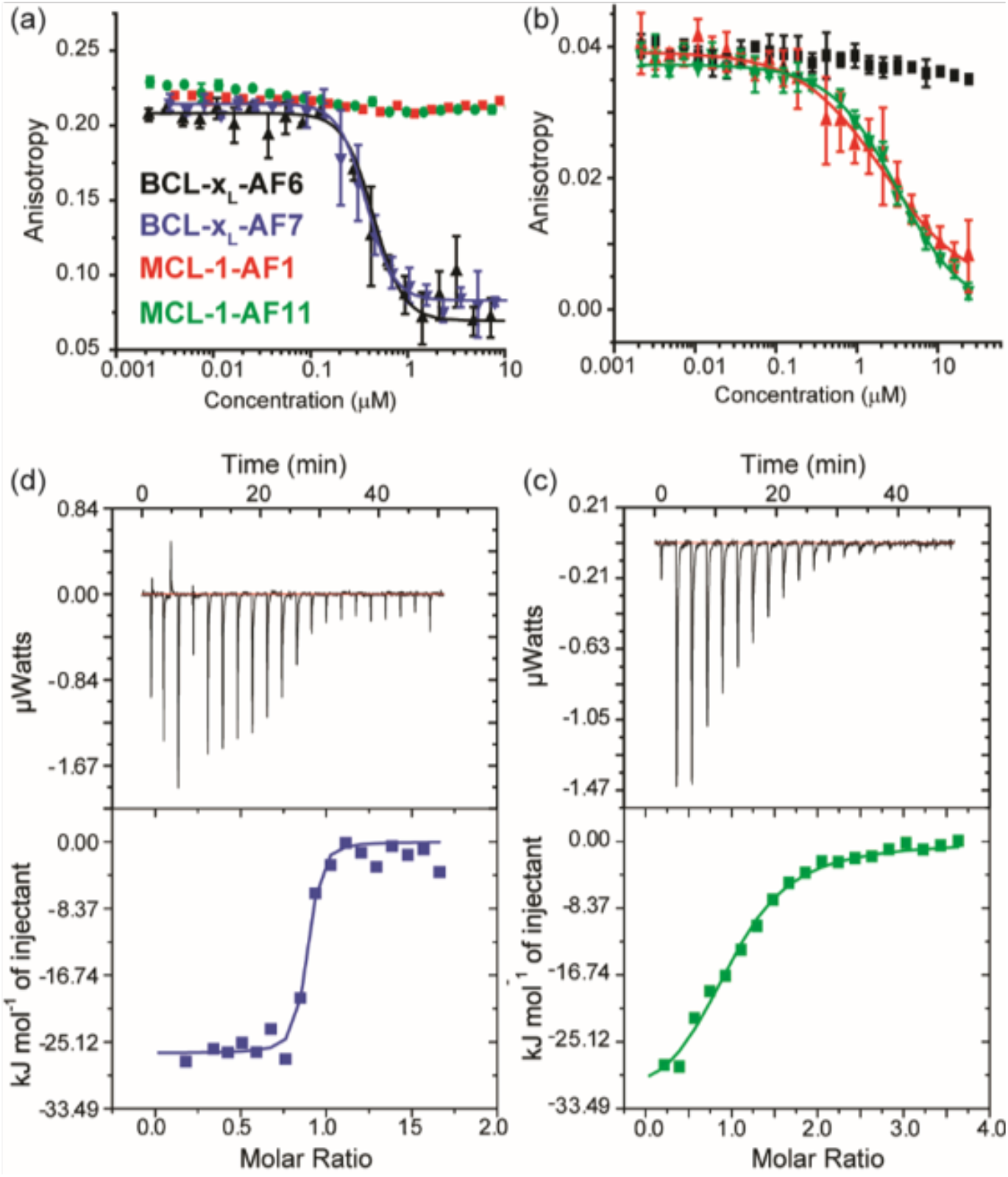
Biophysical Analyses on BCL-x_L_ and MCL-1 binding Affimers. Fluorescence anisotropy competition assays for (a) BCL-x_L_/BODIPY-BAK and (b) MCL-1/FITC-NOXA-B; black = **BCL-x_L_- AF6**, blue = **BCL-x_L_-AF7** red = **MCL-1-AF1**, green = **MCL-1-AF11**. (c-d) ITC data for binding of Affimers to BCL-x_L_ and MCL-1; colour coding as for (a), (c) thermograms and curve fitting for ITC analyses on **BCL-x_L_-AF7/**BCL-x_L_ interaction. (d) thermograms and curve fitting for ITC analyses on **MCL-1-AF11/**MCL-1 interaction.

Isothermal titration calorimetry (ITC; Fig. 3c-d, Fig. S3 and Table 1) confirmed the conclusions garnered from competition FA and gave *K*_d_ values consistent with the determined IC_50_ values. Both **BCL-x_L_-AF6** (BCL-x_L_ selective) and **MCL-1-AF11** (MCL1 selective) Affimers exhibited favourable enthalpic but unfavourable entropic contributions to binding (it was not possible to obtain data for **MCL-AF1**). In the case of **BCL-x_L_-AF6** a particularly strong enthalpic contribution was observed. On the other hand **BCL-x_L_-AF7** was found to be favourable in both the enthalpic and entropic terms. Given the hydrophobic nature of the BH3 binding cleft and high conservation of aliphatic side chains at key positions in both BH3 sequences and the Affimers (see discussion of co-crystal structure below), the observation that different thermodynamic signatures can be employed to achieve recognition could be a useful consideration in informing inhibitor design. Whilst thermodynamic signatures are notoriously difficult to interpret, and enthalpically driven hydrophobic molecular recognition has been documented, the “classical” view of hydrophobic driven binding is one of entropic desolvation.^47–50^ Moreover, our own prior studies characterized BH3/BCL-2 family interactions as entropically driven.^20^

**Table 1.** Thermodynamic parameters of Affimer/BCL-x_L_ and Affimer/MCL-1 binding determined

| Protein | Ligand | K<sub>d</sub> (nM) | ΔH (kJ/mol) | TΔS (J/mol) |
| --- | --- | --- | --- | --- |
| BCL-x<sub>L</sub> | **BCL-x<sub>L</sub>-AF6** | 90.9 ± 3 | -45.3 ± 0.93 | -17.1 |
| BCL-x<sub>L</sub> | **BCL-x<sub>L</sub>-AF7** | 38.6 ± 1 | -26.5 ± 0.75 | 53.0 |
| MCL-1 | **MCL-1-AF11** | 3400 ± 400 | -34.4 ± 1.76 | -11.34 |

Whilst the Affimer technology regularly produces binders with K_d_ in the nanomolar range^36^, here we added multiple layers of screening (inhibition of BH3 binding; compatible with anisotropy and ITC experiments) in addition to panning for high affinity binders. This will naturally lead to an attrition rate where clones that do not meet the criteria are lost, which may explain the slightly lower affinities we observe (Table 1). We have chosen this approach as the selectivity achieved here is of significantly greater value than affinity alone in experiments where inhibition of a single member of a highly homologous family is desired. With additional and more stringent panning and selection (potentially quicker), or a larger library or use of affinity maturation techniques (slower), better affinities may be achievable where all criteria are met.

### Crystal Structures and Conformational Selection

Having established that Affimers act as selective inhibitors of BCL-2 family PPIs, we attempted to obtain co-crystals to allow high-resolution structural interpretation of the interactions. **BCL-x_L_-AF6**/BCL-x_L,_ **BCL-x_L_-AF7**/BCL-x_L_ and **MCL-1-AF11**/MCL-1 co-crystals were obtained (see methods, Table 2 and ESI); and the structures solved by molecular replacement using Phaser.^51^

**Table 2.** Data collection, processing and refinement statistics for Affimer:target complexes

|  | AF6:Bcl-xL | AF7:Bcl-xL | AF11:MCL-1 |
| --- | --- | --- | --- |
| Resolution range (Å) * | 71.8-1.91<br>(1.96-1.91) | 40.39-2.24<br>(2.30-2.24) | 69.96-2.20<br>(2.24-2.20) |
| Space group | *P* 4<sub>2</sub> 2<sub>1</sub> 2 | *P* 2<sub>1</sub> | *C* 2 2 2<sub>1</sub> |
| Unit-cell parameters. a, b, c (Å),<br>α β γ ° | 101.1, 101.1, 102.1,<br>90, 90, 90 | 68.3, 87.3, 112.2,<br>90, 96.2, 90 | 92.14, 107.5, 226.2,<br>90, 90, 90 |
| No. of observed reflections |  | 202,533 |  |
| No. of unique reflections | 41,668 | 61,813 | 56,938 |
| Redundancy |  | 3.3 (3.3) |  |
| Completeness (%) * | 99.88 | 98.1 (97.8) | 99.40 |
| < I/σ(I) > * |  | 12.6 (1.4) |  |
| R<sub>merge</sub> (%) §* |  | 4.1 (83.0) |  |
| R<sub>pim</sub> (%) ¥* |  | 3.9 (78.0) |  |
| CC<sub>1/2</sub> |  | 0.72 |  |
| *R* factor (%) | 22.7 | 23.8 | 26.4 |
| R<sub>free</sub> (%) † | 24.7 | 28.6 | 30.5 |
| No. of protein non-H atoms | 3,677 | 7,402 | 7,478 |
| No. of water molecules | 100 | 0 | 60 |
| R.m.s.d bond lengths (Å) ξ | 0.004 | 0.014 | 0.003 |
| R.m.s.d bond angles (°) ξ | 0.817 | 1.7 | 0.741 |
| Average overall *B* factor (Å<sup>2</sup>) |  |  |  |
| Protein |  | 66.2 |  |
| Residues in the regions of<br>Ramachandran plot (%) ‡ |  |  |  |
| Favoured region | 97.15 | 91.0 | 91.16 |
| Outliers | 0.66 | 2.7 | 6.80 |
| PDB code |  | 6HJL |  |

\*Values given in parentheses correspond to those in the outermost shell of the resolution range.
† R<sub>free</sub> was calculated with 5% of the reflections set aside randomly.
ξ Based on the ideal geometry values of Engh and Huber.
‡ Ramachandran analysis using the program MolProbity [67].

For **BCL-x_L_-AF6**/BCL-x_L,_ the crystals diffracted to 1.91 Å, and the asymmetric unit contains one domain swapped dimer of BCL-x_L_, with one **BCL-x_L_-AF6** bound to the cleft of each monomer (Fig 4 a,c). For **BCL-x_L_-AF7**/BCL-x_L,_ the crystals diffract to 2.24 Å, and the asymmetric unit contains two domain swapped dimers of BCL-x_L_, with one **BCL-x_L_-AF7** bound to the cleft of each monomer (Fig 4 a,b). The residues within the VRs of both Affimers interact with residues lining the BH3-binding cleft on the surface of BCL-x_L_. Representative electron density is presented in Fig. S4. As expected, given that we selected for competitive Affimers, the Affimers bind at the BH3 binding groove. Indeed, the Affimers use some of the available binding pockets in the groove. In **BCL-x_L_-AF6**, F43 binds to the pocket as does F101 on BIM (in PDB 5C3G), and F76 binds the same pocket as I97 on BIM. For **BCL-x_L_- AF7**, W41 binds the same pocket as F101 on BIM. However, the universally conserved Asp to Arg hydrogen-bond is not replicated in any way by the Affimer.

**Figure 4.**
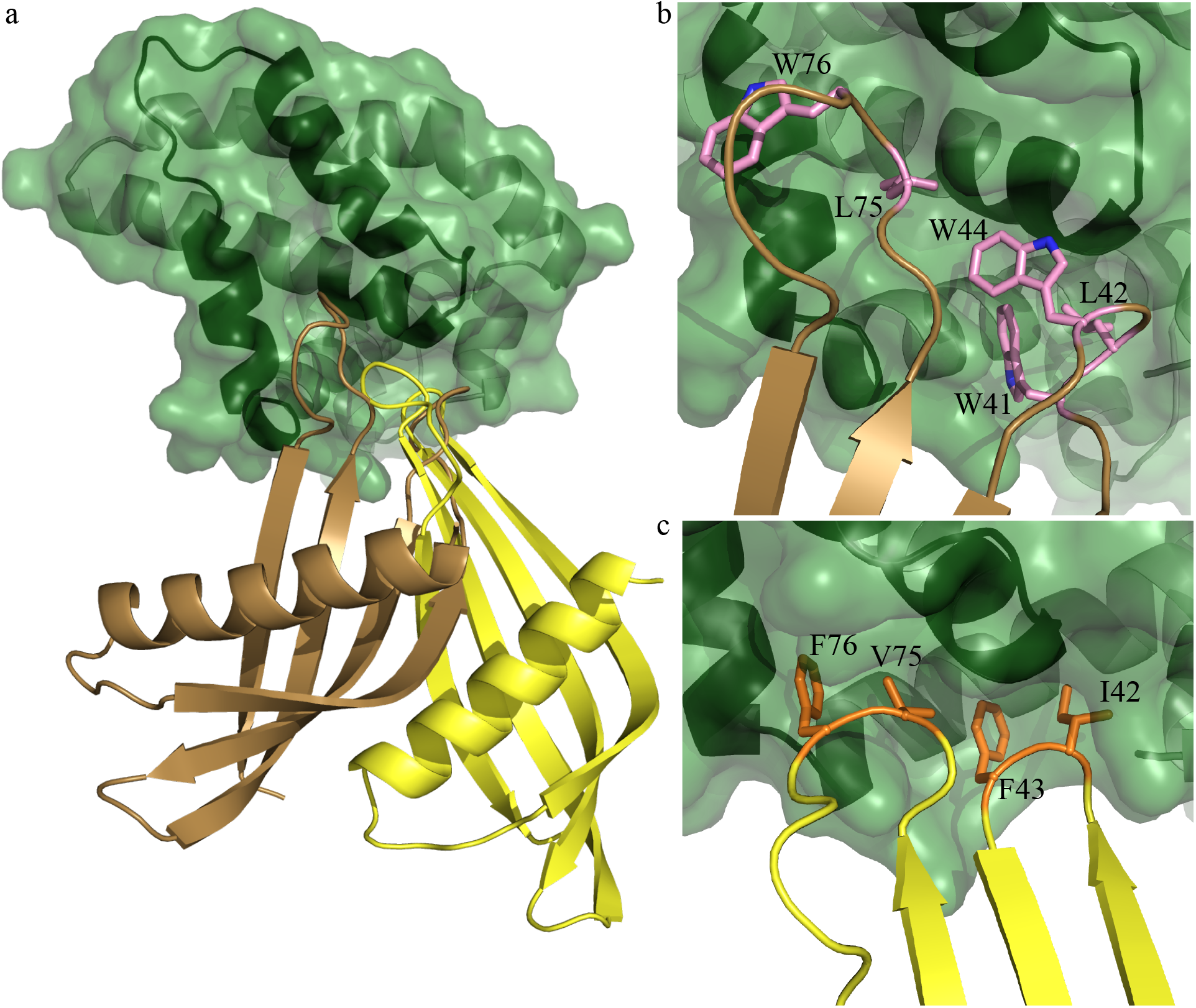
Affimer: BCL-x_L_ co-crystal structures (a) Affimer **BCL-x_L_-AF6** (yellow) or **BCL-x_L_-AF7** (brown) bound to BCL-x_L_ (green) illustrating projection of Affimer loops into BH3 binding cleft (b) zoom in to hydrophobic clusters from Variable Regions 1 & 2 of **AF7** (pink side chains, labelled) projecting into the BH3 (c) hydrophobic cluster from VR 1 & 2 of **AF6** (orange side chains, labelled).

On inspection it is apparent that the Affimers are selecting a single conformation of the BCL-x_L_ domain. The binding groove on BCL-x_L_ is formed by helices 3 and 4 (Fig. 5) and helix 3 is mobile such that the width of the groove can vary. When the BIM BH3 peptide is bound, helix 3 moves to accommodate the peptide in a relatively wide groove. By comparison, when BCL-x_L_ is bound to small molecules such as WEHI-539, or to **BCL-x_L_-AF6** and **BCL-x_L_-AF7**, the groove is narrow (Fig. 5). When bound to BIM peptide the groove is 16.1 Å wide at the widest point (Fig. 5a), whereas in the small molecule and Affimer bound conformation it is 11.0 Å wide (Fig. 5b). All four copies of **BCL-x_L_-AF7**/BCL-x_L_ in the asymmetric unit have this narrow groove i.e. small molecule conformation, suggesting that this is independent of crystal packing. For clarity, an overlay of the BCL-x_L_ domain only, when in complex with BIM, Affimer and WEHI-539, is presented in Fig. S5. This suggests that not only are Affimers selecting a single conformation from the multiple dynamic possibilities in solution, but also that they could be used to select a conformation that is desired by the experimenter. In this case, not only does the Affimer bound conformation correspond to the small molecule bound conformation (thus this might be a better starting point for structure based drug design), but it is also a conformation where the binding groove is too narrow to accommodate the BH3 helix, possibly potentiating the orthosteric competitive effect.

**Figure 5.**
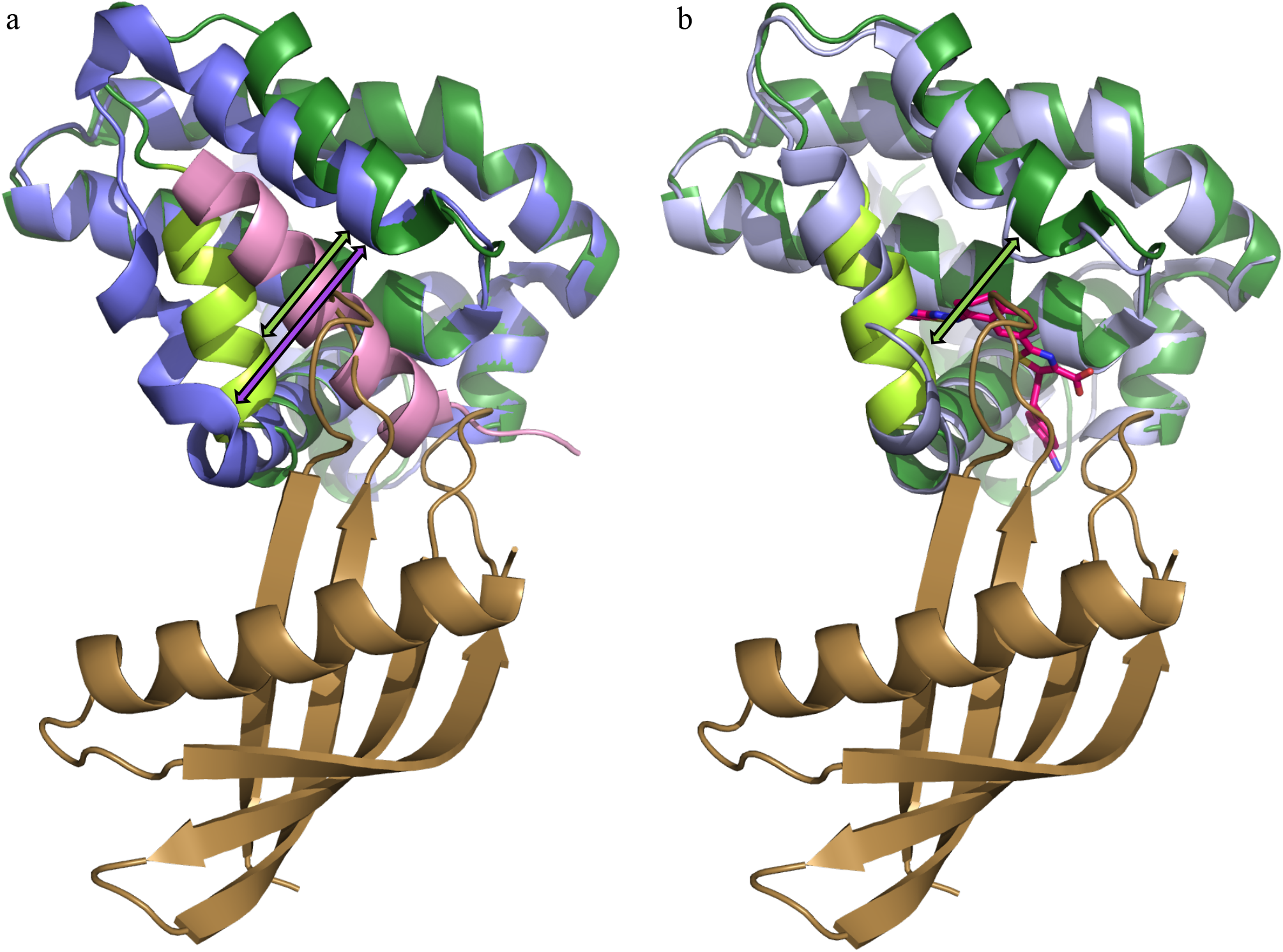
Conformational selection of BCL-x_L_ by the Affimers: When bound to Affimer (**BCL-xL-AF6** or **BCL-xL-AF7**) BCL-x_L_ is in the small molecule bound conformation, not the peptide bound conformation. (a) Comparison between the BCL-xL conformation when bound to **BCL-xL-AF7** (BCL-xL dark green, **BCL-xL-AF7** brown) vs BIM (PDB:5FMK; BCL-x_L_ purple, BIM peptide pink); (b) comparison between the BCL-xL conformation when bound to **BCL-xL-AF7** (dark green) vs **WEHI-539** (PDB:3ZLR; BCL-x_L_ light blue, compound in magenta). Mobile helix 3 of BCL-x_L_ (residues 102-114) is highlighted in light green. Note the position of helix 3: the helix binding groove is 5 Å wider at the last turn when bound to peptide (purple arrow) than to **WEHI-539** or **BCL-xL-AF7** (green arrow; 16.1 Å vs 11.0 Å). This will be a favourable conformation to select for the desired inhibition of peptide binding.

For **MCL-1-AF11**/MCL-1, the crystals diffract to 2.20 Å, and the asymmetric unit contains 4 copies of the complex (Fig 6 a,b) with representative electron density in Fig. S4. Again, the competition with BH3 peptide is mediated via VR residues inserted into the binding groove, with W73 of **MCL-1-AF11** binding in the same pocket as V85 of NOXA (PDB 2NLA); unsurprisingly, all three crystal structures reveal that the Affimers use the available pockets in the peptide binding site for binding. Again, as for the BCL-x_L_ Affimers, we see that the binding of Affimer selects a desirable conformation. Song *et al*.^52^ have shown that MCL-1 can adopt multiple conformations in solution with differing outcomes in the cell. When BIM BH3 is bound, MCL-1 adopts a non-helical conformation at the QRN motif around Arg222. By contrast, when bound to Mule BH3 or a range of small molecules, the QRN motif is helical. Critically, when the QRN motif is helical, then ubiquitin can be added at this motif, and this promotes cellular degradation by the proteasome. By both competitively inhibiting BH3 binding at the groove, and promoting degradation in cells, the small molecules dramatically reduce MCL activity in treated cells, promoting apoptosis in MCL dependent cancer cell lines. Interestingly, our structure shows that **MCL-1-AF11** also selects the desired helical, ubiquitinatable, conformation, again demonstrating that an Affimer can be isolated that selects a specific desired conformation.

**Figure 6.**
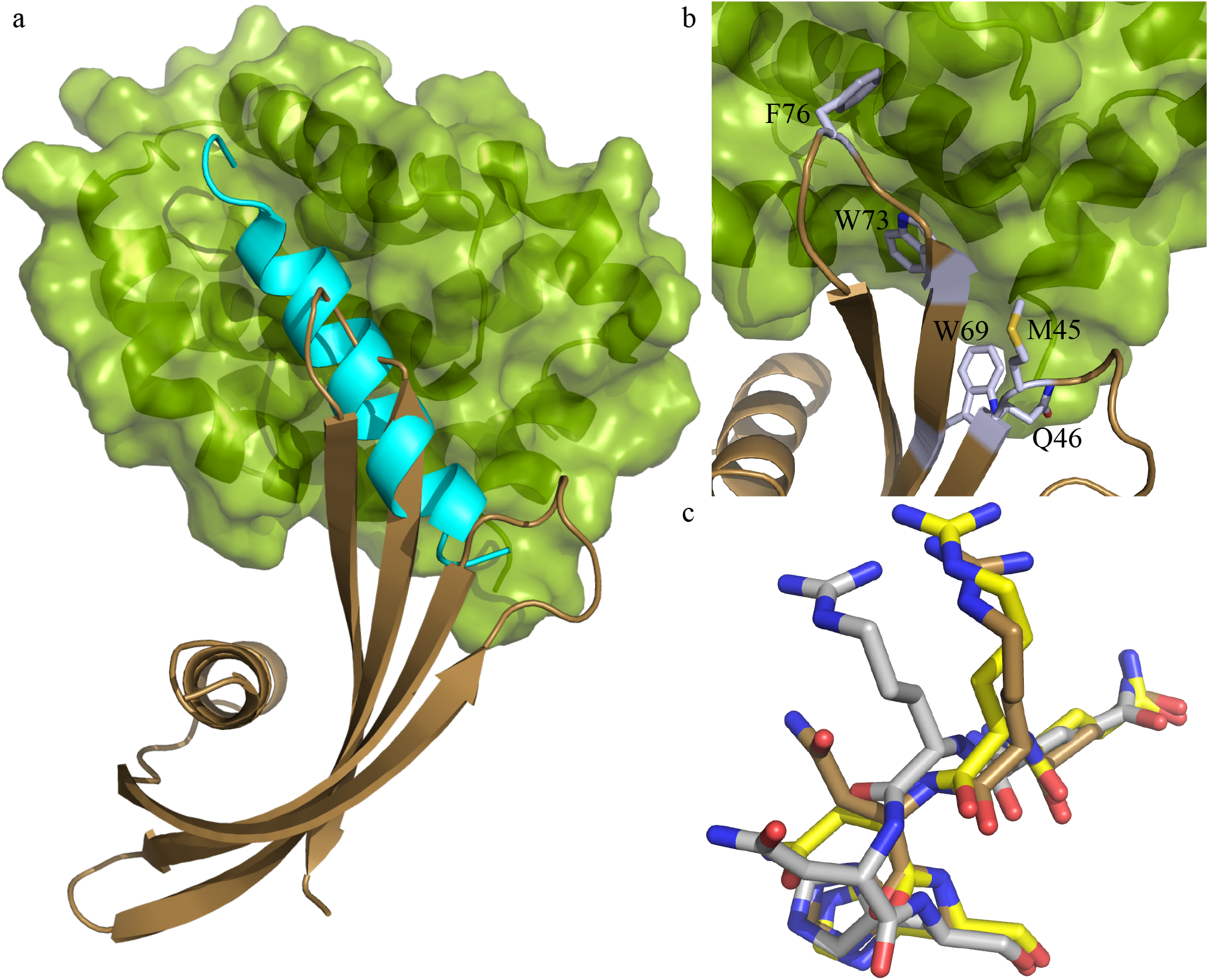
The Affimer/MCL-1 co-crystal structure suggests conformational selection (a) Affimer **MCL-1- AF11** (brown) bound to MCL-1 (green) illustrating projection of Affimer loops into BH3 binding cleft. Overlayed on NOXA (cyan) bound MCL-1 (b) zoom in to hydrophobic clusters from Variable Regions 1 & 2 (grey side chains, labelled) projecting into the BH3 cleft (c) Ubiquitinatable region of MCL-1 bound to BIM (grey), MULE (yellow), and **MCL-1-AF11** (brown). Note the position of the Arginine carbonyl group: Mule and Affimer select the desired ubiquitinatable, degradable conformation, whereas BIM does not.

All known biological partners,^4^ and indeed all designed peptides thus far,^13,14,53–55^ that interact with BCL-2 family proteins at the BH3 groove do so with peptide in a helical conformation. Crucially, the VRs do not adopt an *α*-helical conformation to make interactions with the BH3 binding cleft (presumably in part because they are constrained from doing so). Despite the absence of a helical binding conformation, the Affimers project amino acids side chains so as to mimic key hydrophobic and polar contacts made by BH3 ligands and BCL/MCL. Thus, we have identified proteins that binds to the BH3 binding groove via a non-canonical fold. Peptidomimetics based on this structure, rather than the canonical *α*-helices, may therefore represent a novel starting point for small molecule discovery.

## Discussion

We have used the Affimer libraries to isolate reagents that are highly selective for their targets and can discriminate between very related homologues such as Bcl family proteins and Sumo variants.^35^ Given that Affimers express well in live cells, this approach should prove fruitful in identifying reagents that can be used to investigate cell biology that is dependent on related proteins, for instance signalling pathways where there is a need to discriminate the actions of highly related isoforms.^36^

Comparison between the Affimer/BCL-x_L_ and BCL-x_L_/BIM (PDB ID: 1PQ1) structures illuminates key features; the BH3 cleft narrows in response to Affimer binding in contrast to the wider cleft observed for binding of BIM (Fig. 5a). The BCL-x_L_ conformation in the Affimer bound form is much more similar to that observed for small molecule bound structures such as WEHI-539 (PDB ID: 3ZLR. Fig 5b), where BCL-xL is also domain swapped.

Similarly, when comparing the structures of MCL-1 bound to peptide vs Affimer we observe that the variable loops of the Affimer are inserted into the BH3 binding groove and that a desirable conformation is selected. In this case, the conformation is remote from the binding site, but again reflects a small molecule bound conformation. In contrast to the BIM bound MCL conformation in a non-helical conformation at QRN (around Arg 222) that is not ubiquitinatable, MCL-1 bound to small molecules from Song *et al*.,^52^ UMI-77, Maritoclax and TW-37, or Affimer **MCL-1-AF11**, has a helical conformation around the QRN motif that can be ubiquitinated, providing both orthosteric inhibition of BH3 binding and degradation via the proteasome to potentiate the removal of MCL-1 function in cells. The Affimer again selects a conformation of a target protein to promote a desired functional outcome.

These data, and a previous report,^34^ imply that Affimers can be used not only to identify selective sequences that differentiate between related family members, but also that they can be used for conformational selection of productive or desirable binding modes. The role of conformational selection in studies of protein-protein interactions is increasingly being recognized.^56–61^ Still, it remains a major challenge to account for protein dynamics in structure-based drug-design.^62^ This library of Affimers allows exploration of a dynamic range of protein target conformers, potentially facilitating generation of template pharmacophores for small-molecule ligand design and structure based-ligand design, which may offer an advantage over the current process that typically operates using static crystal structures.^62^ In summary, we have identified non-natural protein ligands that exhibit selectivity for different BCL-2 family members. Although computational protein design has been applied to discovery of BCL-2 family selective binders,^63^ to our knowledge non-antibody based binding proteins have not previously been shown to differentiate between these proteins notably BCL-x_L_ and MCL-1; this is noteworthy given the role of MCL-1 in driving several cancers.^64,65^ We note the attrition rate that is a consequence of applying a variety of activity criteria as we progressed along this pipeline. Of the 12 MCL-1 and 11 BCL-x_L_ binders identified, not all were inhibitors of the cognate BH3 binding, not all were amenable to biophysical assays, and only three have yielded high resolution crystal structures. This serves as a reminder that a large number of binders is required in order to identify ligands with multiple selection criteria applied.

The co-crystal structure provides inspiration for the structure-based design of peptide and small molecule based BCL-2 family modulators, a goal we will pursue in due course. Similarly, BCL-2 binding Affimers themselves could be elaborated for therapeutic or diagnostic use.^34,66^ Of potentially equal significance is the observation that non-canonical folds can substitute for native folds in peptide/protein based inhibitors of PPIs ^67,68^; this is reminiscent of the use of a β-hairpin to mimic an α-helix for p53/*h*DM2 inhibition.^69,70^ In contrast to those studies, the sequences identified here were obtained under selection pressure, and this poses the question: do BCL-2 family proteins function in cells through molecular modes of interaction other than the canonical α-helix/cleft motif observed to date?

## Methods

### Overexpression and Purification of BCL-2 Family proteinxs

BCL-2 family proteins were expressed and purified following previously published methods^20^ – a full description is given in the supporting information.

### Screening for Affimers

BCL-2 family proteins were biotinylated using EZ-link NHS-SS-biotin (Pierce), according to the manufacturer’s instructions. Biotinylation was confirmed using streptavidin conjugated to horseradish peroxidase (HRP). Biotin-BCL-2 family proteins were added and incubated on pre-blocked steptavdin plate, the plate was then washed using a KingFisher robotic platform (ThermoFisher) and 10^12^ cfu of the prepanned phage library was added and incubated for 2.5 h with shaking. Wells were wash ten times and eluted with 100 µL 0.2 M glycine (pH 2.2) for ten minutes neutralized with 15 µL 1 M Tris-HCl (pH 9.1), further eluted with triethylamine 100 mM for 6 min, and neutralised with 1 M Tris–HCl (pH 7). Eluted phage were used to infect ER2738 cells for 1 h at 37 °C and 90 rpm then plated onto LB agar plates with 100 µg/ml carbenicillin and grown overnight. All colonies were scrapped into 5 mL of 2XYT with carbenicillin (10 µg/mL) and 1 x 10^9^ M13K07 helper phage were added. After an overnight incubation phage were precipitated with 4 % polyethylene glycol 8000, 0.3 M NaCl and resuspended in 1 ml of 10 mM Tris, pH 8.0, 1 mM EDTA (TE buffer). 2 µL phage suspension was used for the second round panning round using streptavidin magnetic beads as opposed to streptavidin plates (Invitrogen); otherwise the second pan was conducted in the same way as the first pan. The third pan was conducting using neutravidin high binding capacity plates (Pierce). After the final pan colonies were picked, an ELISA was conducted to select positive clones (in the same way as the enrichment ELISA) which were sent for Sanger sequencing.

### Overexpression and purification of Affimers

The Affimers were subcloned from the phage display vector into pET11a then expressed and purified from *E. coli* strain Rosetta 2. 10 ml of overnight starter culture was used to inoculate 1 L 2 x YT containing 125 μg/ml Ampicillin Cultures were grown at 37 °C until OD600 ~ 0.6 – 0.8, the temperature was then switched to 18 °C and protein expression induced by the addition of 0.5mM IPTG. Induced cultures were grown at 18 °C overnight before harvesting by centrifugation (Beckman JLS 8.100 rotor, 4,500 rpm, 12 min, 4 °C). Cells were resuspended in 50 mM TRIS pH 8.0, 500 mM NaCl, 15 mM imidazole and lysed by sonication in the presence of 10 μL of 1 U.ml-1 DNase I per litre of over-expression culture and cell lysate was clarified (Beckman JA25.50 rotor, 17,000 rpm, 45 min, 4 °C). The supernatant was filtered (0.45 μM syringe filter) before application onto a 5 ml HisTrap that had previously been equilibrated with 50 mM TRIS pH 8.0, 500 mM NaCl, 15 mM imidazole. The cleared cell lysate was then allowed to flow through the HisTrap with the aid of a peristaltic pump. The HisTrap was then washed with 10 CV of 50 mM TRIS pH 8.0, 500 mM NaCl, 15 mM imidazole followed by 10 CV 50 mM TRIS pH 8.0, 500 mM NaCl, 50 mM imidazole and 10 CV 50 mM TRIS pH 8.0, 500 mM NaCl, 100 mM imidazole. The Affimer was then eluted from the HisTrap with 50 mM TRIS pH 8.0, 500 mM NaCl, 300 mM imidazole. Successful elution was confirmed on a gel before further purification was undertaken. The eluted Affimer was concentrated (Amicon Ultra centrifugal filter, MWCO 10,000) to approximately 5 ml. The sample was then filtered before being loaded onto a Superdex 75 column (GE healthcare) equilibrated in 50 mM TRIS pH 8.0, 250 mM NaCl, 0.5 mM DTT, 2.5% Glycerol. The protein eluted as a monomer from gel filtration. The purified protein was concentrated to ~ 6 mg/ml and stored at – 80 °C with the addition of 5% Glycerol. Additionally, Affimers **BCL-x_L_-AF6** and **MCL-1-AF11** were subcloned into pET28a His-SUMO expression vector to remove flexible residues at the *N* and *C*-termini which have hindered crystallisation. The constructs were over-expressed in the *E. coli* strain Rosetta 2. 10 ml of overnight starter culture was used to inoculate 1 L 2 xYT containing 50 μg/ml Kanamycin. Cultures were grown at 37 °C until OD600 ~ 0.6 – 0.8, the temperature was then switched to 18 °C and protein expression induced by the addition of 0.5uM IPTG. Induced cultures were grown at 18 °C overnight before harvesting by centrifugation (Beckman JLS 8.100 rotor, 4,500 rpm, 12 min, 4 °C). Cells were re-suspended in 20 mM TRIS pH 8.0, 500 mM NaCl, 15 mM imidazole and lysed by sonication and cell lysate was clarified (Sorvall SS34 rotor, 17,000 rpm, 45 min, 4 °C). The supernatant was filtered (0.45 μM syringe filter) before application onto a 5 ml HisTrap that had previously been equilibrated with 20 mM TRIS pH 8.0, 500 mM NaCl, 15 mM imidazole. The HisTrap was then washed with 10 CV of 50 mM TRIS pH 8.0, 500 mM NaCl, 15 mM imidazole followed by 10 CV 50 mM TRIS pH 8.0, 500 mM NaCl, 50 mM imidazole. The His-SUMO-Affimer fusion protein was then eluted from the HisTrap with 20 mM TRIS pH 8.0, 500 mM NaCl, 300 mM imidazole. The His-SUMO-Affimer fusion protein was cleaved overnight in dialysis into 20 mM TRIS pH 8.0, 250 mM NaCl in the presence of Smt3 protease, Ulp1, overnight at 4 °C. To remove any uncleaved Affimer, His-SUMO and Ulp1, the sample was reapplied to a HisTrap in 20 mM TRIS pH 8.0, 250 mM NaCl and the flow through containing Affimer collected. This was concentrated (Amicon Ultra centrifugal filter, MWCO 10,000) to approximately 10 ml. The sample was then filtered before being loaded onto a Superdex 75 column (GE healthcare) equilibrated in 20 mM TRIS pH 8.0, 250 mM NaCl, 0.5 mM DTT, 2.5% Glycerol. The protein eluted as a monomer from gel filtration. The purified protein was concentrated to ~ 6 mg/ml and stored at – 80 °C with the addition of 5% Glycerol.

### Isothermal Titration Calorimetry

Kinetic information of the Affimer interactions with the Bcl-2 family proteins was established using the ITC200 microcalorimeter (MicroCal, Northampton, MA). Sample buffers were 50 mm Tris, pH 7.9, 200 mM NaCl. Experiments were carried out at 30 °C. The syringe contained 40 μl of 350 μM Affimer (**BCL-x_L_-AF6**, **BCL-x_L_-AF7** or **MCL-1-AF11**); 2 μl injections were applied every 180 seconds. The cell contained 205 μl of 35 μM BCL-x_L_ or MCL-1. Cell concentrations were adjusted to a 1:1 stoichiometric interaction and Microcal Origin software version 7.0 was used to determine the dissociation constants (*K*_d_). All measurements were repeated at least twice.

### Single Point Fluorescence Anisotropy

A single point assay was carried out at a fixed concentration of Affimer (1 µM), FITC-NOXA-B or BODIPY-BAK at 87.5nM or 37.5nM respectively and MCL-1 or BCL-x_L_ at 175 nM or 50nM respectively in phosphate buffer (40 mM sodium phosphate, 200 mM sodium chloride, 0.02 mg ml^-1^ Bovine serum albumin, pH 7.50). Each Affimer was assessed in triplicate and left to equilibrate for 45 minutes in the dark. A positive control was present on each test plate (BAK peptide for the BCL-x_L_/BAK interaction and NOXA-B peptide for the MCL-1/NOXA-B interaction) at the same concentration as the test compounds. Anisotropy values were then determined and a percentage of efficiency was calculated for each compound relative to BAK or NOXA-B’s efficiency, with blank wells set to zero.

### Competition assays

Competition fluorescence anisotropy assays and data processing were performed adapting previously described protocols.^20^ Briefly, the buffer used for fluorescence anisotropy was phosphate buffer (40 mM sodium phosphate, 200 mM sodium chloride, 0.02 mg ml^-1^ Bovine serum albumin, pH 7.50). Assays were run in 384 well Optiplates and were scanned using a Perkin Elmer EnVisionTM 2103 MultiLabel plate reader. Fluorescein labelled peptides used an excitation and emission wavelength of 490 nm and 535 nm respectively whilst BODIPY labelled peptides used an excitation and emission wavelength of 531 nm and 595 nm respectively, with a bandwidth of 5 nm. BODIPY-BAK/BCL-x_L_ competition assays were performed in 384 well plates in phosphate buffer with the concentration of the inhibitor typically starting from 5-50 µM, diluted over 24 points in a 2/3 regime with [BODIPY-BAK] and [BCL-x_L_] fixed at 50 nM and 150 nM respectively. Plates were read after 45 minutes of incubation. FITC-NOXA-B/MCL-1 competition assays were performed in a 384 well plates in phosphate buffer, with the concentration of the inhibitor typically starting from 5-100 µM, diluted over 24 points in a 2/3 regime and with [FITC-NOXA-B] and [MCL-1] fixed at 50 nM and 150 nM respectively. Plates were read after 45 minutes.

### Co-crystallisation

BCL-x_L_ was incubated with an excess of **BCL-x_L_-AF7** overnight, before co-purification in 200mM NaCl, 50mM TRIS pH 8.0, 0.5mM DTT via gel filtration on a Superdex75 column. Crystals grew in 12% PEG 1500, 0.1M Sodium Acetate pH 5.5, 2.5 M NaCl, 1.5% MPD at 20°C at 5mg/ml using the sitting drop vapour diffusion method. The crystals were cryoprotected in 20% glycerol and data collected at the Diamond Light Source on beamline i04-1 to 2.24 Å resolution at 100K. The diffraction images were integrated, scaled and reduced using the suite of program XIA2^71^ with five percent of the reflections selected at random and excluded from the refinement using FREERFLAG.^72^ The unit cell parameters for the crystal are a=68.3Å, b=87.3Å, c=112.2Å, α=90.0°, β=96.2°, γ=90.0° in space group P21 with four **BCL-x_L_-AF7**/BCL-_XL_ complexes in the asymmetric unit cell. The data processing statistics are shown in Table 2. The structure was determined by molecular replacement using the program PHASER^51^ with the human BCL-_XL_ structure (PDB code 1R2D),^73^) and the truncated Affimer (PDB code 4N6U,^39^) as the search models. Manual inspection of electron density maps with iterative cycles of model building and refinement were carried out using COOT^74^ and REFMAC5.^75,76^ During the course of model building structural validations were carried out using the program MOLPROBITY.^77^ All refinement statistics are shown in Table 2. The structures have been deposited in the Protein Data Bank (www.pdb.org) and has been assigned the PDB codes xxxx (**BCL-x_L_-AF6**/Bcl-_xL_) 6HJL (**BCL-x_L_-AF7**/Bcl-_xL_) and xxxx **MCL-1-AF11**/MCL-1). Structures were analysed and figures prepared with PyMol^78^.

BCL-x_L_ was incubated with an excess of **BCL-x_L_-AF6** at room temperature for 3 hours. The complex was purified via gel filtration on a Superdex 200 column equilibrated with 20 mM Tris pH 7.4, 50 mM NaCl, 2 mM DTT. Crystals grew in 0.1 M Tris pH 7, 0.2 M MgCl2, 10% w/v PEG 8K at 20°C at 10 mg ml^-1^ concentration using the sitting drop vapour diffusion method. The crystals were cryoprotected with 20% glycerol and data was collected on beamline i04-1 at the Diamond Light Source to 1.90 Å. The unit cell parameters for the crystal are a=101.1Å, b=101.1Å, c=102.1Å, α=90.0°, β=90°, γ=90.0° in space group P42 21 2 with two **BCL-x_L_-AF6**/BCL-x_L_ complexes in the asymmetric unit cell.

The structure was solved using similar strategy as detailed above for **BCL-x_L_-AF7**/BCL-x_L_ complex.

MCL-1 was incubated with an excess of **MCL-1-AF11** at room temperature for 3 hours. The complex was purified via gel filtration on a Superdex 200 column equilibrated with 20 mM Tris pH 7.4, 50 mM NaCl, 2 mM DTT. Crystals grew in 0.1 M Na Acetate pH 4.6, 30 % w/v PEG MME 2K, 0.2 M (NH_4_)_2_SO_4_ at 20°C at 9 mg ml^-1^ concentration using the sitting drop vapour diffusion method. The crystals were cryoprotected with 20% glycerol and data was collected on beamline i04-1 at the Diamond Light Source to 2.21 Å.

The unit cell parameters for the crystal are a=92.14Å, b=107.5Å, c=226.2Å, α=90.0°, β=90°, γ=90.0° in space group C 2 2 21 with four **MCL-1-AF11**/MCL-1 complexes in the asymmetric unit cell.

## Acknowledgements

We thank Pallavi Ramsahye for protein purification, Dr Nasir Khan for assistance with CD and Iain Manfield for help with ITC. We acknowledge Diamond Light Source for time on MX beamlines under proposal MX10305 and thank the beamline scientists especially at beamline i04-1. This work was supported by the EPSRC (EP/N013573/1), the ERC (ERC-StG-240324) and The Wellcome Trust (097827/Z/11/A, WT094232MA, 094232/Z/10/Z)

## Author Contributions

All authors have read and approved the manuscript. FH and JAM performed experiments, analysed data and wrote the manuscript. JT, PRR, HFK, CT, CHT, BJ, FN and BIMW performed experiments and analysed data. DCT and JC designed experiments and applied for funding. AJW and TAE applied for funding, designed and performed experiments, analysed data and wrote the manuscript

## Additional Information

The authors declare no competing interests.

## References

1. Milroy, L.-G., Grossmann, T.N., Hennig, S., Brunsveld, L. & Ottmann, C. Modulators of Protein–Protein Interactions. Chemical Reviews 114, 4695–4748 (2014).

2. Arkin, M.R., Tang, Y. & Wells, J.A. Small-Molecule Inhibitors of Protein-Protein Interactions: Progressing toward the Reality. Chemistry & Biology 21, 1102–1114 (2014).

3. Chen, H.-C. et al. An interconnected hierarchical model of cell death regulation by the BCL-2 family. Nat Cell Biol 17, 1270–1281 (2015).

4. Kvansakul, M. & Hinds, M. The Bcl-2 family: structures, interactions and targets for drug discovery. Apoptosis 20, 136–150 (2015).

5. Volkmann, N., Marassi, F.M., Newmeyer, D.D. & Hanein, D. The rheostat in the membrane: BCL-2 family proteins and apoptosis. Cell Death Differ 21, 206–215 (2014).

6. Vela, L. & Marzo, I. Bcl-2 family of proteins as drug targets for cancer chemotherapy: the long way of BH3 mimetics from bench to bedside. Current Opinion in Pharmacology 23, 74–81 (2015).

7. Lessene, G., Czabotar, P.E. & Colman, P.M. BCL-2 family antagonists for cancer therapy. Nat Rev Drug Discov 7, 989–1000 (2008).

8. Czabotar, P.E., Lessene, G., Strasser, A. & Adams, J.M. Control of apoptosis by the BCL-2 protein family: implications for physiology and therapy. Nat Rev Mol Cell Biol 15, 49–63 (2014).

9. Certo, M. et al. Mitochondria primed by death signals determine cellular addiction to antiapoptotic BCL-2 family members. Cancer cell 9, 351–365 (2006).

10. Chen, L. et al. Differential Targeting of Prosurvival Bcl-2 Proteins by Their BH3-Only Ligands Allows Complementary Apoptotic Function. Molecular cell 17, 393–403 (2005).

11. Pelay-Gimeno, M., Glas, A., Koch, O. & Grossmann, T.N. Structure-Based Design of Inhibitors of Protein–Protein Interactions: Mimicking Peptide Binding Epitopes. Angewandte Chemie International Edition 54, 8896–8927 (2015).

12. Azzarito, V., Long, K., Murphy, N.S. & Wilson, A.J. Inhibition of [alpha]-helix-mediated protein-protein interactions using designed molecules. Nature Chemistry 5, 161–173 (2013).

13. Foight, G.W., Ryan, J.A., Gullá, S.V., Letai, A. & Keating, A.E. Designed BH3 Peptides with High Affinity and Specificity for Targeting Mcl-1 in Cells. ACS Chemical Biology 9, 1962–1968 (2014).

14. London, N., Gullá, S., Keating, A.E. & Schueler-Furman, O. In Silico and in Vitro Elucidation of BH3 Binding Specificity toward Bcl-2. Biochemistry 51, 5841–5850 (2012).

15. Chen, T.S., Palacios, H. & Keating, A.E. Structure-Based Redesign of the Binding Specificity of Anti-Apoptotic Bcl-xL. Journal of Molecular Biology 425, 171–185 (2013).

16. DeBartolo, J., Dutta, S., Reich, L. & Keating, A.E. Predictive Bcl-2 Family Binding Models Rooted in Experiment or Structure. Journal of Molecular Biology 422, 124–144 (2012).

17. Walensky, L.D. & Bird, G.H. Hydrocarbon-Stapled Peptides: Principles, Practice, and Progress. Journal of Medicinal Chemistry 57, 6275–6288 (2014).

18. LaBelle, J.L. et al. A stapled BIM peptide overcomes apoptotic resistance in hematologic cancers. Journal of Clinical Investigation 122, 2018–2031 (2012).

19. Cromm, P.M., Spiegel, J. & Grossmann, T.N. Hydrocarbon Stapled Peptides as Modulators of Biological Function. ACS Chemical Biology 10, 1362–1375 (2015).

20. Miles, J.A. et al. Hydrocarbon constrained peptides - understanding preorganisation and binding affinity. Chemical Science 7, 3694–3702 (2016).

21. Stewart, M.L., Fire, E., Keating, A.E. & Walensky, L.D. The MCL-1 BH3 helix is an exclusive MCL-1 inhibitor and apoptosis sensitizer. Nat Chem Biol 6, 595–601 (2010).

22. Walensky, L.D. et al. A Stapled BID BH3 Helix Directly Binds and Activates BAX. Molecular cell 24, 199–210 (2006).

23. Walensky, L.D. et al. Activation of Apoptosis in Vivo by a Hydrocarbon-Stapled BH3 Helix. Science 305, 1466–1470 (2004).

24. Barnard, A. et al. Selective and Potent Proteomimetic Inhibitors of Intracellular Protein– Protein Interactions. Angewandte Chemie International Edition 54, 2960–2965 (2015).

25. Azzarito, V. et al. Stereocontrolled Protein Surface Recognition Using Chiral Oligoamide Proteomimetic Foldamers. Chemical Science 6, 2434–2443 (2015).

26. Checco, J.W. et al. α/β-Peptide Foldamers Targeting Intracellular Protein–Protein Interactions with Activity in Living Cells. Journal of the American Chemical Society (2015).

27. Smith, B.J. et al. Structure-Guided Rational Design of α/β-Peptide Foldamers with High Affinity for BCL-2 Family Prosurvival Proteins. ChemBioChem 14, 1564–1572 (2013).

28. Leverson, J.D. et al. Potent and selective small-molecule MCL-1 inhibitors demonstrate on-target cancer cell killing activity as single agents and in combination with ABT-263 (navitoclax). Cell Death Dis 6, e1590 (2015).

29. Souers, A.J. et al. ABT-199, a potent and selective BCL-2 inhibitor, achieves antitumor activity while sparing platelets. Nat Med 19, 202–208 (2013).

30. Wilson, W.H. et al. Navitoclax, a targeted high-affinity inhibitor of BCL-2, in lymphoid malignancies: a phase 1 dose-escalation study of safety, pharmacokinetics, pharmacodynamics, and antitumour activity. The Lancet Oncology 11, 1149–1159 (2010).

31. Aguilar, A. et al. A Potent and Highly Efficacious Bcl-2/Bcl-xL Inhibitor. Journal of Medicinal Chemistry 56, 3048–3067 (2013).

32. Gemperli, A.C., Rutledge, S.E., Maranda, A. & Schepartz, A. Paralog-Selective Ligands for Bcl-2 Proteins. Journal of the American Chemical Society 127, 1596–1597 (2005).

33. Chin, J.W. & Schepartz, A. Design and Evolution of a Miniature Bcl-2 Binding Protein. Angewandte Chemie International Edition 40, 3806–3809 (2001).

34. Robinson, J.I. et al. Affimer proteins inhibit immune complex binding to FcγRIIIa with high specificity through competitive and allosteric modes of action. Proceedings of the National Academy of Sciences of the United States of America 115, E72–E81 (2018).

35. Hughes, D.J. et al. Generation of specific inhibitors of SUMO-1– and SUMO-2/3–mediated protein-protein interactions using Affimer (Adhiron) technology. Science Signaling 10, 10.1126/scisignal.aaj2005 (2017).

36. Tiede, C. et al. Affimer proteins are versatile and renewable affinity reagents. eLife 6, e24903 (2017).

37. Arrata, I., Barnard, A., Tomlinson, D.C. & Wilson, A.J. Interfacing native and non-native peptides: using Affimers to recognise [small alpha]-helix mimicking foldamers. Chemical Communications 53, 2834–2837 (2017).

38. Kyle, H.F. et al. Exploration of the HIF-1α/p300 interface using peptide and Adhiron phage display technologies. Molecular BioSystems 11, 2738–2749 (2015).

39. Tiede, C. et al. Adhiron: a stable and versatile peptide display scaffold for molecular recognition applications. Protein Engineering Design and Selection 27, 145–155 (2014).

40. Nord, K., Nilsson, J., Nilsson, B., Uhlén, M. & Nygren, P.-Å. A combinatorial library of an α- helical bacterial receptor domain. Protein Engineering Design and Selection 8, 601–608 (1995).

41. Koide, A., Bailey, C.W., Huang, X. & Koide, S. The fibronectin type III domain as a scaffold for novel binding proteins11Edited by J. Wells. Journal of Molecular Biology 284, 1141–1151 (1998).

42. Binz, H.K., Stumpp, M.T., Forrer, P., Amstutz, P. & Plückthun, A. Designing Repeat Proteins: Well-expressed, Soluble and Stable Proteins from Combinatorial Libraries of Consensus Ankyrin Repeat Proteins. Journal of Molecular Biology 332, 489–503 (2003).

43. Koide, A., Wojcik, J., Gilbreth, R.N., Hoey, R.J. & Koide, S. Teaching an Old Scaffold New Tricks: Monobodies Constructed Using Alternative Surfaces of the FN3 Scaffold. Journal of Molecular Biology 415, 393–405 (2012).

44. Škrlec, K., Štrukelj, B. & Berlec, A. Non-immunoglobulin scaffolds: a focus on their targets. Trends in Biotechnology 33, 408–418 (2015).

45. Woodman, R., Yeh, J.T.H., Laurenson, S. & Ferrigno, P.K. Design and Validation of a Neutral Protein Scaffold for the Presentation of Peptide Aptamers. Journal of Molecular Biology 352, 1118–1133 (2005).

46. Kyle, H.F. et al. Exploration of the HIF-1[small alpha]/p300 interface using peptide and Adhiron phage display technologies. Molecular BioSystems 11, 2738–2749 (2015).

47. Biela, A. et al. Dissecting the Hydrophobic Effect on the Molecular Level: The Role of Water, Enthalpy, and Entropy in Ligand Binding to Thermolysin. Angewandte Chemie International Edition 52, 1822–1828 (2013).

48. Snyder, P.W. et al. Mechanism of the hydrophobic effect in the biomolecular recognition of arylsulfonamides by carbonic anhydrase. Proceedings of the National Academy of Sciences 108, 17889–17894 (2011).

49. Setny, P., Baron, R. & McCammon, J.A. How Can Hydrophobic Association Be Enthalpy Driven? Journal of Chemical Theory and Computation 6, 2866–2871 (2010).

50. Schauperl, M., Podewitz, M., Waldner, B.J. & Liedl, K.R. Enthalpic and Entropic Contributions to Hydrophobicity. Journal of Chemical Theory and Computation 12, 4600–4610 (2016).

51. McCoy, A.J., et al. Phaser crystallographic software. Journal of Applied Crystallography 40, 658–674 (2007).

52. Song, T. et al. Deactivation of Mcl-1 by Dual-Function Small-Molecule Inhibitors Targeting the Bcl-2 Homology 3 Domain and Facilitating Mcl-1 Ubiquitination. Angewandte Chemie International Edition 55, 14250–14256 (2016).

53. Frappier, V., Jenson, J.M., Zhou, J., Grigoryan, G. & Keating, A.E. Tertiary Structural Motif Sequence Statistics Enable Facile Prediction and Design of Peptides that Bind Anti-apoptotic Bfl-1 and Mcl-1. Structure 27, 606–617.e5 (2019).

54. Jenson, J.M. et al. Peptide design by optimization on a data-parameterized protein interaction landscape. Proceedings of the National Academy of Sciences (2018).

55. Dutta, S., Chen, T.S. & Keating, A.E. Peptide Ligands for Pro-survival Protein Bfl-1 from Computationally Guided Library Screening. ACS Chemical Biology 8, 778–788 (2013).

56. Berlow, R.B., Dyson, H.J. & Wright, P.E. Hypersensitive termination of the hypoxic response by a disordered protein switch. Nature 543, 447–451 (2017).

57. Rogers, J.M. et al. Interplay between partner and ligand facilitates the folding and binding of an intrinsically disordered protein. Proceedings of the National Academy of Sciences 111, 15420–15425 (2014).

58. Bah, A. et al. Folding of an intrinsically disordered protein by phosphorylation as a regulatory switch. Nature 519, 106–109 (2015).

59. Staus, D.P. et al. Allosteric nanobodies reveal the dynamic range and diverse mechanisms of G-protein-coupled receptor activation. Nature 535, 448 (2016).

60. Irannejad, R. et al. Conformational biosensors reveal GPCR signalling from endosomes. Nature 495, 534 (2013).

61. Steyaert, J. & Kobilka, B.K. Nanobody stabilization of G protein-coupled receptor conformational states. Current Opinion in Structural Biology 21, 567–572 (2011).

62. Śledź, P. & Caflisch, A. Protein structure-based drug design: from docking to molecular dynamics. Current Opinion in Structural Biology 48, 93–102 (2018).

63. Berger, S. et al. Computationally designed high specificity inhibitors delineate the roles of BCL2 family proteins in cancer. eLife 5, e20352 (2016).

64. Nhu, D., Lessene, G., Huang, D.C.S. & Burns, C.J. Small molecules targeting Mcl-1: the search for a silver bullet in cancer therapy. MedChemComm 7, 778–787 (2016).

65. Varadarajan, S. et al. Evaluation and critical assessment of putative MCL-1 inhibitors. Cell Death Differ 20, 1475–1484 (2013).

66. Xie, C. et al. Development of an Affimer-antibody combined immunological diagnosis kit for glypican-3. Scientific Reports 7, 9608 (2017).

67. Wuo, M.G. & Arora, P.S. Engineered protein scaffolds as leads for synthetic inhibitors of protein–protein interactions. Current Opinion in Chemical Biology 44, 16–22 (2018).

68. Sha, F., Salzman, G., Gupta, A. & Koide, S. Monobodies and other synthetic binding proteins for expanding protein science. Protein Science 26, 910–924 (2017).

69. Fasan, R. et al. Structure–Activity Studies in a Family of β-Hairpin Protein Epitope Mimetic Inhibitors of the p53–HDM2 Protein–Protein Interaction. ChemBioChem 7, 515–526 (2006).

70. Fasan, R. et al. Using a beta-hairpin to mimic an alpha-helix: Cyclic peptidomimetic inhibitors of the p53-HDM2 protein-protein interaction. Angewandte Chemie-International Edition 43, 2109–2112 (2004).

71. Winter, G. xia2: an expert system for macromolecular crystallography data reduction. Journal of Applied Crystallography 43, 186–190 (2010).

72. Brünger, A.T. [19] Free R value: Cross-validation in crystallography. in Methods in Enzymology, Vol. 277 366–396 (Academic Press, 1997).

73. Manion, M.K. et al. Bcl-XL Mutations Suppress Cellular Sensitivity to Antimycin A. Journal of Biological Chemistry 279, 2159–2165 (2004).

74. Emsley, P., Lohkamp, B., Scott, W.G. & Cowtan, K. Features and development of Coot. Acta Crystallographica Section D 66, 486–501 (2010).

75. Murshudov, G.N., Vagin, A.A. & Dodson, E.J. Refinement of Macromolecular Structures by the Maximum-Likelihood Method. Acta Crystallographica Section D 53, 240–255 (1997).

76. Vagin, A.A. et al. REFMAC5 dictionary: organization of prior chemical knowledge and guidelines for its use. Acta Crystallographica Section D 60, 2184–2195 (2004).

77. Chen, V.B. et al. MolProbity: all-atom structure validation for macromolecular crystallography. Acta Crystallographica Section D 66, 12–21 (2010).

78. The PyMOL Molecular Graphics System, Version 2.0 Schrödinger, LLC.

