## Supporting Information for "Selective Affimers Recognize BCL-2 Family Proteins Through Non-Canonical Structural Motifs"

**Additional Materials and Methods**

### BCL-$x_l$ and MCL-1 expression

BCL-$x_l$ and MCL-1 were overexpressed and purified according to our published procedure.[1] The pet28a His-SUMO Mcl-1 (172-327) and pet28a His-SUMO Bcl-$x_l$ (a chimera with BCL-2, 1-198, missing 26-81) constructs were over-expressed in the *E.coli* strain Rosetta 2. 10 ml of overnight starter culture was used to inoculate 1 L 2 xYT containing 50 µg/ml Kanamycin. Cultures were grown at 37 °C until OD600 ~ 0.6 – 0.8, the temperature was then switched to 18 °C and protein expression induced by the addition of 0.5uM IPTG. Induced cultures were grown at 18 °C overnight before harvesting by centrifugation (Beckman JLS 8.100 rotor, 4,500 rpm, 12 min, 4 °C). Cells were resuspended in 20mM TRIS pH 8.0, 500mM NaCl, 15mM Imidazole and lysed by sonication and cell lysate was clarified (Sorvall SS34 rotor, 17,000 rpm, 45 min, 4 °C). The clarified lysate was applied to a 5ml HisTrap equilibrated with 20mM TRIS pH 8.0, 500mM NaCl, 15mM Imidazole. The HisTrap was then washed with 10 CV of 20mM TRIS pH 8.0, 500mM NaCl, 15mM Imidazole followed by 10 CV 20mM TRIS pH 8.0, 500mM NaCl, 25mM Imidazole. The fusion proteins were eluted in 20mM TRIS pH 8.0, 500mM NaCl, 300mM Imidazole. The His-SUMO tag was removed in overnight in dialysis into 20mM TRIS pH 8.0, 250mM NaCl in the presence of Smt3 protease, Ulp1, overnight at 4°C. Uncleaved material and His-SUMO were removed by reapplication of the sample to a HisTrap in 20mM TRIS pH 8.0, 250mM NaCl and the flow through containing Mcl-1 or Bcl-xl collected. The proteins were then filtered before further purification on a Superdex 75 (GE healthcare) equilibrated in 20mM TRIS pH 8.0, 250mM NaCl, 0.5mM DTT, 2.5% Glycerol. Purified proteins were concentrated and stored at -80°C.

### BCL-2 expression

Glutathione S-transferase (GST) tagged BCL-$2^{1-205}$ fusion protein constructs were over-expressed in *E.coli* BL21 (DE3) Rosetta 2, and purified. 10 ml of overnight starter culture was used to inoculate 1 L 2 xYT containing 100 µg/ml Ampicillin. Cultures were grown at 37 °C until $OD_{600}$ reached 0.6 – 0.8. The temperature was reduced to 18 °C and protein expression induced by adding 0.3 mM IPTG. Induced cultures were grown at 18°C overnight before harvesting by centrifugation (Beckmann JLS 8.1, 5000rpm, 15 minutes, 4°C). Cell pellets were re-suspended in lysis buffer (50mM TRIS pH 8.0, 200mM NaCl, 5mM DTT, 1mM EDTA) and lysed by sonication in the presence of 10 µL of 1 U.ml-1 DNase I per litre of over-expression culture. Cell lysate was clarified (Sorvall SS34 rotor, 18,000 rpm, 45 min, 4°C), and the supernatant was filtered (0.45 µM Minisart, Sartorius). Clarified cell lysate was added to approximately 10 mL Glutathione Superflow Resin (Generon) packed in a free-flow gravity column. The column was washed with 5 column volumes of water and equilibrated with 5 column volumes of lysis buffer. The lysate was then added to the column and placed on an analogue roller mixer (SKS science) at 4°C for 1-3 hours. Lysate was then eluted from the column under gravity, and the resin was washed first with 100 mL high salt wash buffer (50 mM Tris pH 8, 1 mM DTT, 1 mM EDTA, 500 mM NaCl, 10% glycerol, 0.01% Triton), then 100 mL low salt wash buffer (50 mM Tris pH 8, 1 mM DTT, 1 mM EDTA, 200 mM NaCl, 10% glycerol, 0.01% Triton). The resin was re-suspended in 20 mL of low salt wash buffer, supplemented with 400 µL PreScission protease to cleave the GST tag from the fusion protein. Following overnight incubation on an analogue roller mixer (SKS science) at 4°C, cleaved protein was obtained by collecting the flow through from the column. To collect all cleaved protein, the resin was washed with 50 mL of low salt wash buffer, and the flow through collected. All wash fractions were collected and analysed by SDS-PAGE. Cleaved protein was concentrated (Amicon, MWCO 10,000) to 5 ml, filtered (0.22 µM Minisart, Sartorius) and further purified by Size Exclusion Chromatography. Purified protein was concentrated and stored at -80°C, with the addition of 5% glycerol to aid long term stability.

**BAK and BAX expression**

Mxe intein / chitin binding domain (CBD) tagged fusion protein constructs were over-expressed in *E.coli* C41 (DE3) cells, and purified. 2 mL of overnight starter culture was used to inoculate 1 L LB containing 100 µg/mL Ampicillin. Cultures were grown at 37°C until $OD_{600}$ reached 0.6 – 0.8. For $BAX^{1\text{-}171(C62S\ C126S)}$ protein expression was induced by adding 0.1 mM IPTG. Induced cultures were grown at 28°C overnight. For $BAK^{16\text{-}185(C166S)}$ protein expression was induced by adding 1 mM IPTG, and induced cultures were grown at 37°C for 4 hours. Cells were harvested by centrifugation (Beckmann JLS 8.1, 5000rpm, 15minutes, 8°C). Cell pellets were resuspended in lysis buffer (20 mM HEPES pH7.0, 100 mM NaCl, 1 mM EDTA)) and lysed by sonication. Cell lysate was clarified (Sorvall SS34 rotor, 18,000 rpm, 45 min, 8°C), the supernatant was filtered (0.22 µM Minisart, Sartorius). Clarified cell lysate was added to approximately 20 mL Chitin Resin (New England Biolabs) packed in a free-flow gravity column, equilibriated with 20 column volumes of lysis buffer. The lysate was then added to the column and eluted from the column at a flow rate of 1 mL/min, then the resin was washed with 10 column volumes of lysis buffer. The column was quickly equilibrated with 3 column volumes of lysis buffer supplemented with 50 mM DTT. The resin was then resuspended in 1 column volume of lysis buffer supplemented with 50 mM DTT. Following overnight incubation on an analogue roller mixer (SKS science) at 25°C, cleaved protein was obtained by collecting the flow through from the column. To collect all cleaved protein, the resin was washed with 50 mL of lysis buffer, then a further 100 mL of lysis buffer. All wash fractions were collected and analysed by SDS-PAGE. Columns were stored in 20% ethanol. Cleaved protein was concentrated (Amicon, MWCO 10,000) to 10 mL, filtered (0.22 μM Minisart, Sartorius) and further purified by Size Exclusion Chromatography. Purified protein was stored at 4°C without further concentration.

### Additional Data and Figures

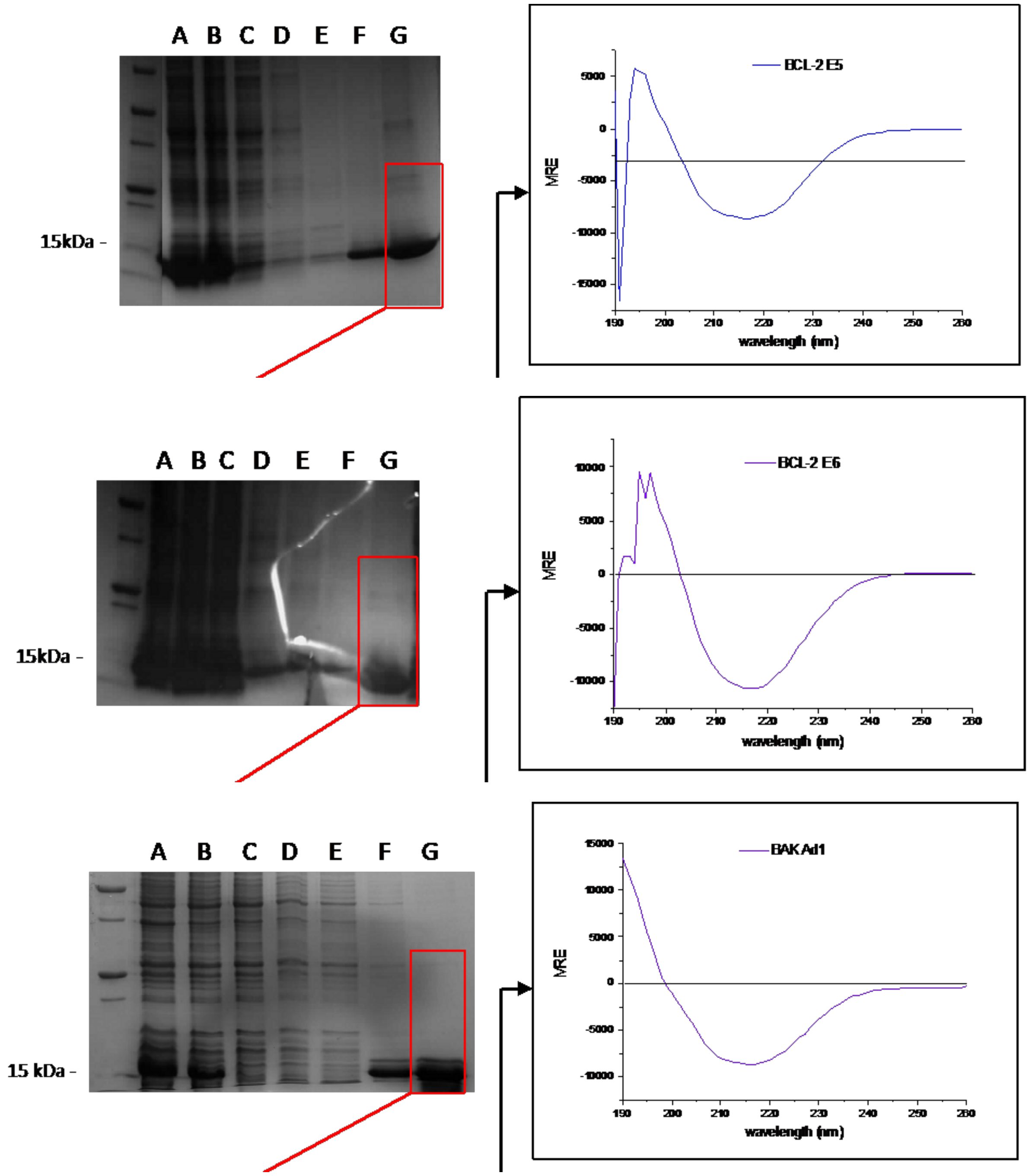


**Figure S1**. Purification and representative CD analyses for BCL-2 and BAK binding Affimers

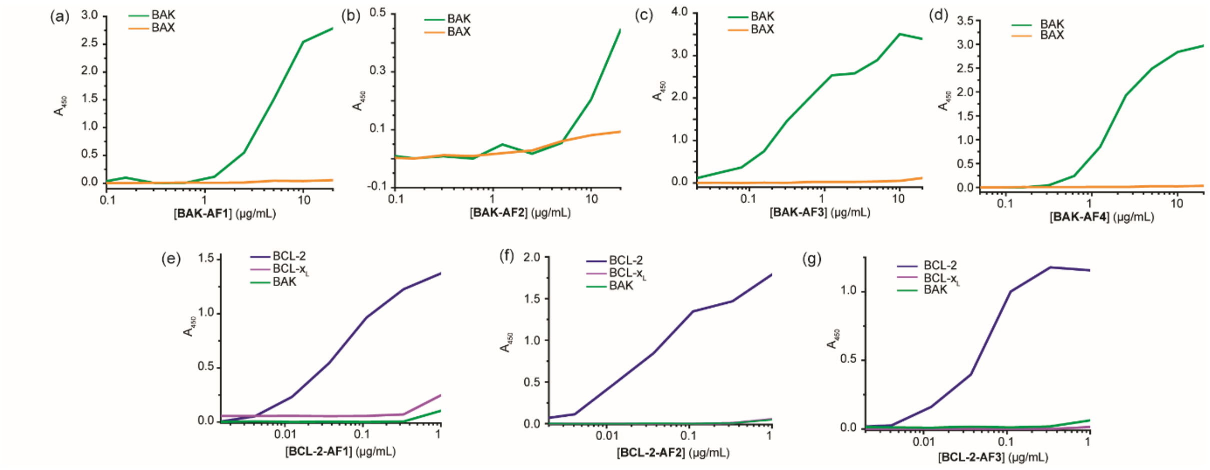


**Figure S2**. Binding analyses of BCL-2 family binding Affimers; binding ELISA for (a) **BAK-AF1**, (b) **BAK-AF2**, (c) **BAK-AF3**, (d) **BAK-AF4**, (e) **BCL-2-AF3**, (f) **BCL-2-AF3**, (g) **BCL-2-AF3**.

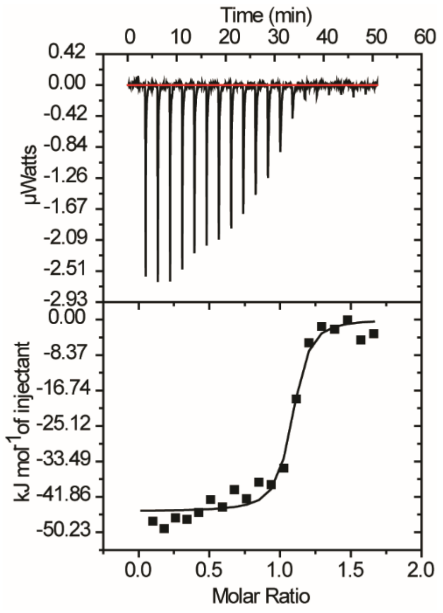


**Figure S3**. ITC data for Binding of **BCL-x$_L$-AF6** to BCL-x$_L$

**Table S1.** sequence information and frequency for Affimers selected against BCL-2 Family

| Protein | Ligand | VR1 | VR2 | Frequency |
|---|---|---|---|---|
| MCL-1 | **MCL-1-AF1** | T T P E P Y E Q Q | W W Q G F Q G M V | 5 |
| MCL-1 | **MCL-1-AF2** | W D Y M G Y S D D | W W Y G F T G W Q | 2 |
| MCL-1 | **MCL-1-AF3** | S R E N V E S W A | F W Q G F F S I M | 7 |
| MCL-1 | **MCL-1-AF4** | F S D T P A Q D S | E W M G F F S M A | |
| MCL-1 | **MCL-1-AF5** | K A Q K E V V A E | F W Q G F Y N W V | 2 |
| MCL-1 | **MCL-1-AF6** | K S A Y D G A W D | N W M G F Y N W D | |
| MCL-1 | **MCL-1-AF7** | W R M Q Y R I G W | P V Y F S N P A M | |
| MCL-1 | **MCL-1-AF8** | V F P S Q D P Q Q | Y W M G F I S W A | |
| MCL-1 | **MCL-1-AF9** | T P F A G D Q M Q | T W K G F F K N D | |
| MCL-1 | **MCL-1-AF10** | F P W M D W L D Q | W W Q G F F Q V E | |
| MCL-1 | **MCL-1-AF11** | M G V N P E E M Q | W W W G F H I W D | |
| MCL-1 | **MCL-1-AF12** | S R E N V E S W A | F W Q G F F S I M | |
| BCL-$x_L$ | **BCL-$x_L$-AF1** | Q F G M A W Y H S | E F V N Q P W S S | 2 |
| BCL-$x_L$ | **BCL-$x_L$-AF2** | H A R D D C M V L | T F H I A G Y I S | 6 |
| BCL-$x_L$ | **BCL-$x_L$-AF3** | Q T Q L W L S L V | N L D Q R R G L M | 6 |
| BCL-$x_L$ | **BCL-$x_L$-AF4** | S V Y M G I S V L | L L K V L D I W H | 2 |
| BCL-$x_L$ | **BCL-$x_L$-AF5** | Y E I G Y T K N S | V L W N V I Y K V | |
| BCL-$x_L$ | **BCL-$x_L$-AF6** | E R N S I F E E F | R D L V F G G P E | |
| BCL-$x_L$ | **BCL-$x_L$-AF7** | M F S W L D W E E | P A L L W S P H G | |
| BCL-$x_L$ | **BCL-$x_L$-AF8** | E Q R L W F S N V | L S G A R R G I Y | |
| BCL-$x_L$ | **BCL-$x_L$-AF9** | M T Q Y D A Y R N | D I P G L F V N L | |
| BCL-$x_L$ | **BCL-$x_L$-AF10** | Q H H W F E F V D | Q D F F D Q Y R P | |
| BCL-$x_L$ | **BCL-$x_L$-AF11** | S Y E V W A F R Q | R D D Y F W L P K | |
| BCL-2 | **BCL-2-AF1** | S A F S W W E R I | D H S Q P I Q M R | 9 |
| BCL-2 | **BCL-2-AF2** | G I G F W V A F S | S S K P N Q M M M | |
| BCL-2 | **BCL-2-AF3** | F M E N N G F V M | K W F F L P P I N | 9 |
| BCL-2 | **BCL-2-AF4** | K M H N S H I I M | K W Y L F P I G A | |

**Table S2.** sequence information and frequency for Affimers selected against BCL-2 Family

| Protein | Ligand | VR1 | VR2 | Frequency |
|---|---|---|---|---|
| BAX | **BAX-AF1** | H V Q A H F W S I | P T E N I M G L K | 22 |
| BAX | **BAX-AF2** | M Q L S S T R L W | T K Y T I Y N P I | |
| BAX | **BAX-AF3** | E E A V P W Y M P | N Y I K L K W H H | |
| BAX | **BAX-AF4** | Q V H M H W Y H A | H L K Q H P L I K | |
| BAX | **BAX-AF5** | N Q A P T F F Q Y | E T K W S H F I Q | |
| BAK | **BAK-AF1** | V G R H S D W A P | Y P I Q S H P P W | 11 |
| BAK | **BAK-AF2** | F V F P H D Y F P | E I A Q W N G W F | 2 |
| BAK | **BAK-AF3** | I M A R P P F E I | A Q L K K T L Y W | 8 |
| BAK | **BAK-AF4** | I W N N S H Y I P | Y P L S L K P F I | |

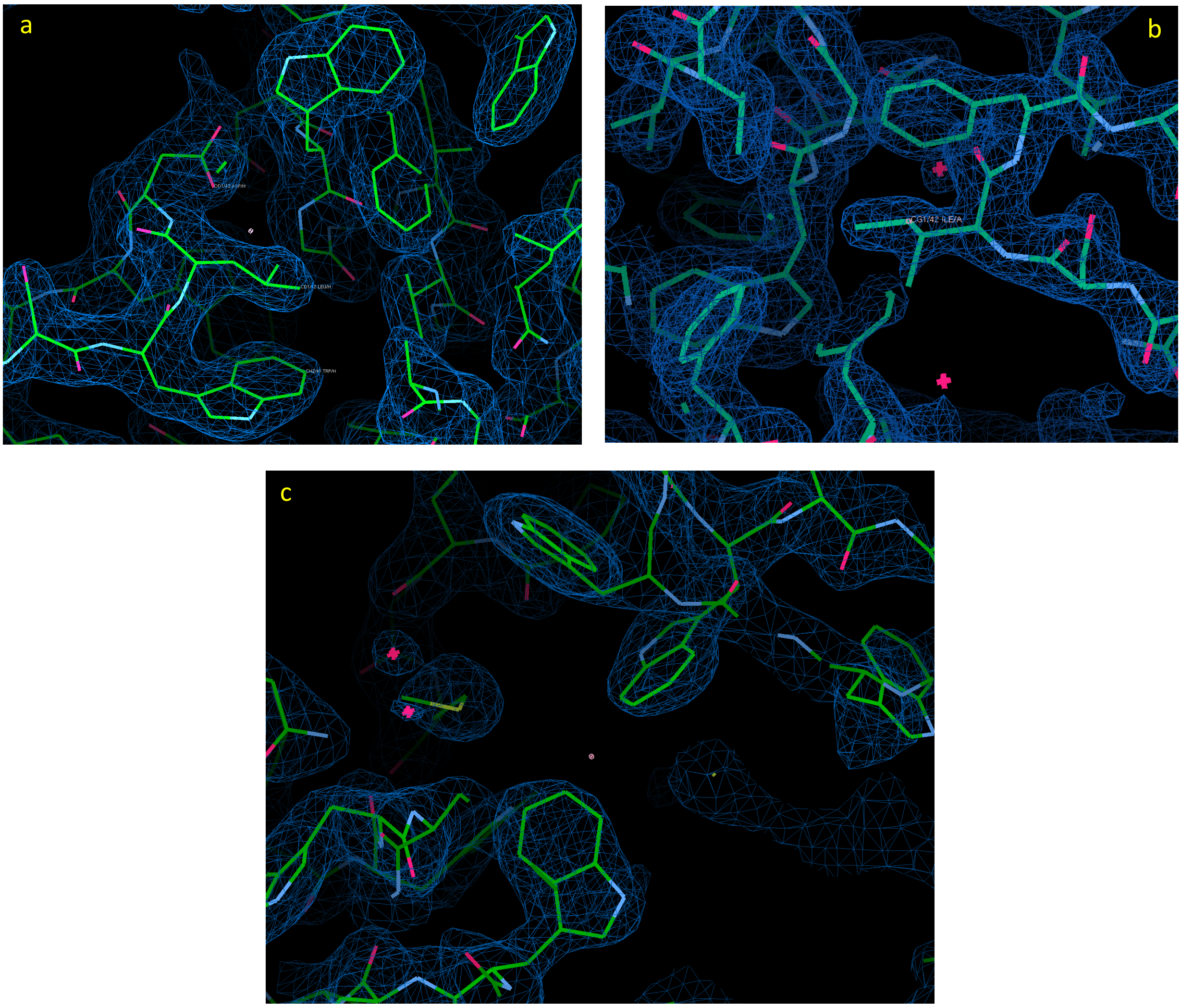


**Figure S4.** Sample 2Fo-Fc electron density at the final stage of refinement for crystal structures. (a) The interface between Affimer **BCL-x$_L$-AF7** and BCL-xL; (b) interface between Affimer **BCL-x$_L$-AF6** and BCL-xL; (c) interface between Affimer **MCL-AF11** and MCL-1. Despite slightly higher than usual $R_{work}$ and $R_{free}$ (for **BCL-x$_L$-AF7** and **MCL-1-AF11,** probably because of a high overall Wilson B-factor for this crystal), the electron density is absolutely unambiguous. Figure generated using *Coot*[2].

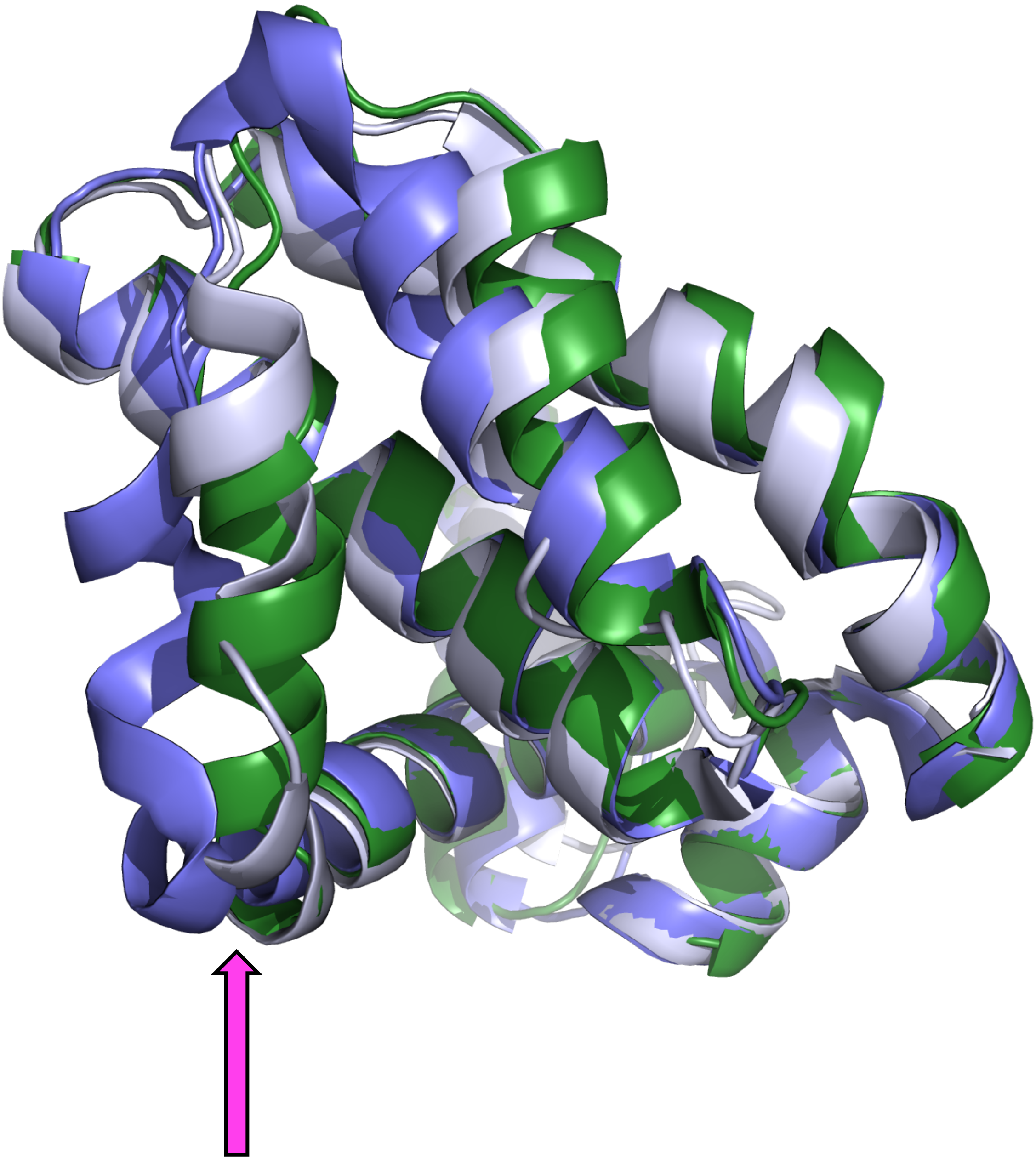

**Figure S5.** Overlay of Bcl-xL when bound in different complexes. Overlay of Bcl-xL when bound to Affimer (dark green) BIM peptide (purple) or WEHI-539 (light blue). Note the position of the BCL-xL helix to the left of peptide/compound (arrow): the helix binding groove is wider when bound to peptide than to WEHI-539 or BCL-xL-AF7.

### References


1. Miles, J. A., Yeo, D. J., Rowell, P., Rodriguez-Marin, S., Pask, C. M., Warriner, S. L., Edwards, T. A. & Wilson, A. J. Hydrocarbon constrained peptides - understanding preorganisation and binding affinity. *Chem. Sci.* **7**, 3694-3702, (2016).

Collaborative Computational Project, Number 4. 1994.

"The CCP4 Suite: Programs for Protein Crystallography". Acta Cryst. D50, 760-763
